# Gonadal androgens are associated with decreased type I interferon production by pDCs and increased IgG titres to BNT162b2 following co-vaccination with live attenuated influenza vaccine in adolescents

**DOI:** 10.1101/2023.08.01.551423

**Authors:** Oliver Sampson, Cecilia Jay, Emily Adland, Anna Csala, Nicholas Lim, Stella M Ebbrecht, Lorna C Gilligan, Angela E Taylor, Sherley S George, Stephanie Longet, Lucy C Jones, Ellie Barnes, John Frater, Paul Klenerman, Susie Dunachie, Miles Carrol, James Hawley, Wiebke Arlt, Andreas Groll, Philip Goulder

## Abstract

mRNA vaccine technologies introduced following the SARS-CoV-2 pandemic have highlighted the need to better understand the interaction of adjuvants and the early innate immune response. Interferon type I (IFN-I) is an integral part of this early innate response and can prime several components of the adaptive immune response. Females are widely reported to respond better than males to seasonal tri- and quad-valent influenza vaccines. Plasmacytoid dendritic cells (pDCs) are the primary cell type responsible for IFN-I production and female pDCs produce more IFN-I than male pDCs since the upstream receptor TLR7 is encoded by the X-chromosome and is biallelically expressed by up to 30% of female immune cells. Additionally, the TLR7 promoter contains putative androgen response elements and androgens have been reported to suppress pDC IFN-I *in-vitro*.

Unexpectedly, therefore, we recently observed that male adolescents mount stronger antibody responses to the Pfizer BNT162b2 mRNA vaccine than female adolescents after controlling for natural SARS-CoV-2 infection. We here examined pDC behaviour in this cohort to determine the impact of IFN-I on anti-Spike and anti-receptor-binding domain titres to BNT162b2. Through LASSO modelling we determined that serum free testosterone was associated with reduced pDC IFN-I but, contrary to the well-described immunosuppressive role for androgens, the more potent androgen dihydrotestosterone was associated with increased IgG titres to BNT162b2. Also unexpectedly, we observed that co-vaccination with live-attenuated influenza vaccine boosted the magnitude of IgG responses to BNT162b2. Together these data support a model where systemic IFN-I increased vaccine-mediated immune responses, but for vaccines with intracellular stages, modulation of the local IFN-I response may alter antigen longevity and consequently vaccine-driven immunity.

**Author Summary:** Type I interferons (IFN-I) are potent antiviral proteins which play a central role in activating the immune response and driving inflammation. IFN-I is predominantly produced by plasmacytoid dendritic cells (pDCs) and female pDCs produce more IFN-I than male pDCs. Consequently, females typically generate stronger antibody responses to vaccines such as seasonal influenza vaccines. In addition, females typically suffer more serious adverse events from vaccines. However, we recently reported in a study of adolescents that males generate stronger antibody responses to the SARS-CoV-2 mRNA vaccine BNT162b2 than females. Here we examine the IFN-I response of pDCs in adolescents co-/vaccinated with BNT162b2 and live-attenuated influenza vaccine (LAIV). We find that male sex hormones reduce pDC IFN-I but are associated with increased BNT162b2 antibody titres. We also observe that LAIV boosts BNT162b2 antibody titres through possible bystander activation of immune cells. These findings are consistent with a reportedly higher incidence of adverse events among males associated with this vaccine. Together these data suggest that IFN-I production typically enhances vaccine-specific immune responses but for new mRNA vaccines such as BNT162b2, that are modified to reduce innate immunogenicity, localised dampening of the IFN-I response in vaccinated tissue by male sex hormones may further delay the clearance of the vaccine, increasing vaccine antigen exposure and allowing time for a stronger antibody response.

## Introduction

The emergence of mRNA vaccine technologies in response to the SARS-CoV-2 pandemic has exposed the gap in understanding the complex interaction of vaccine adjuvants and the innate immune response in shaping the adaptive immune response and outcome to vaccination. Historic understanding of vaccine immunology is based largely on protein-subunit vaccines where empirically-proven adjuvants are proposed to activate antigen presenting cells to present vaccine antigen to CD4+ T-cells, CD8+ T-cells, and B-cells for a robust adaptive response [1,2]. Interferon Type I (IFN-I) is a family of innate antiviral proteins with powerful immunostimulatory effects which include promoting development of CD4+ and CD8+ T-cells to effector and memory phenotypes, and aiding B-cell differentiation and survival [3].

Accordingly, IFN-I is implicated in vaccine outcome and a high initial IFN-I response correlates with increased antibody titres for both the seasonal trivalent influenza vaccine (TIV) and the live-attenuated influenza vaccine (LAIV) in adults [4,5] and more importantly also in children [6,7].

Further, a stronger antibody response to seasonal TIV is consistently reported for females despite a new formulation annually [8,9,10,11,12]. Mechanistically females produce greater quantities of IFN-I since the upstream receptor, TLR7, is encoded by the X-chromosome and escapes X-chromosome inactivation in up to 30% of female immune cells [13]. The primary route of IFN-I induction by TIV is via TLR7 and vaccine-driven IFN-I was shown to induce CD4+ T-cells and B-cells which were essential for protection from lethal influenza challenge in mice [14].

Plasmacytoid dendritic cells (pDCs) are the primary source of IFN-I and pDCs from females consistently produce greater quantities of IFN-I in response to TLR7 stimulation when compared to male pDCs [15,16,17,18,19,20,21,22].

Additionally, male androgens are broadly considered to suppress the immune response [11] and a systems analysis of seasonal TIV responses identified a negative relationship between TIV antibody titres and serum testosterone in adult males [10,12,23]. Indeed, putative androgen response elements are located within the promoters of both TLR7 and the downstream MyD88 signalling intermediate [12,24], providing one mechanism by which androgens may dampen the pDC IFN-I response.

Accordingly, female pDCs treated *in-vitro* with dihydrotestosterone (DHT), the more bioactive product of testosterone, show reduced IFN-I production following TLR7 stimulation [25]. Further, pDCs from male infants during the so-called “mini-puberty” of early infancy – when androgen levels transiently rise to pubertal levels – show reduced IFN-I production in response to TLR7 stimulation [25].

The immune system of children differs markedly from adults and healthy children have higher IFN-I responses than healthy adults, including to natural SARS-CoV-2 infection which has been linked with their increased capacity to clear the virus [26]. Further, a stronger early IFN-I response was predictive of increased neutralising antibody titres following Oxford/AstraZeneca ChAdOx1-nCoV19 (AZD1222) vaccination [27].

In Autumn 2021 adolescent children in the UK were offered a primary dose of Pfizer BNT162b2 alongside intranasal LAIV as part of the nationwide governmental pandemic response. A key immunological feature of BNT162b2 is an early IFN-I response [28] but no difference in outcome to vaccination between sexes has been reported in studies in adults [29,30,31]. Likewise, LAIV was approved for use in children in the UK in 2013 but we are aware of no study to date reporting differences in outcome between sexes in children or adults.

We have recently published that, contrary to seasonal influenza vaccine responses, the anti-Spike and anti-receptor-binding domain (RBD) IgG titres were stronger for males than females in adolescents co-vaccinated with BNT162b2 and LAIV [32]. In this study we characterised the pDC response in 33 adolescents aged 12yrs to 16yrs from this cohort receiving BNT162b2/LAIV co-/vaccination to investigate whether differential IFN-I production drives a differential outcome to co-/vaccination in adolescents.

## Results

### Cohort Demographics

In Autumn 2021 adolescent school children in the United Kingdom were offered a primary dose of Pfizer BNT162b2 alongside optional co-administered nasal LAIV. We enrolled 33 adolescents from three schools in Oxfordshire and their demographics are detailed in Table 1. Baseline blood samples were taken immediately following co-/vaccination (“V1”) to determine prior SARS-CoV-2 infection, androgen levels, and to assess the pDC response to TLR7 stimulation at baseline. The pDC response was also measured on the day after vaccination for a subset (n=18) to determine the influence of co-/vaccination on the early IFN-I response by pDCs. All 33 adolescents were re-sampled 5 weeks post-vaccination (median 37 days; henceforth “post-V1”) to determine the IgG response to the key antigens for BNT162b2: SARS-CoV-2 Spike and receptor-binding domain (RBD).

**Table 1–.**
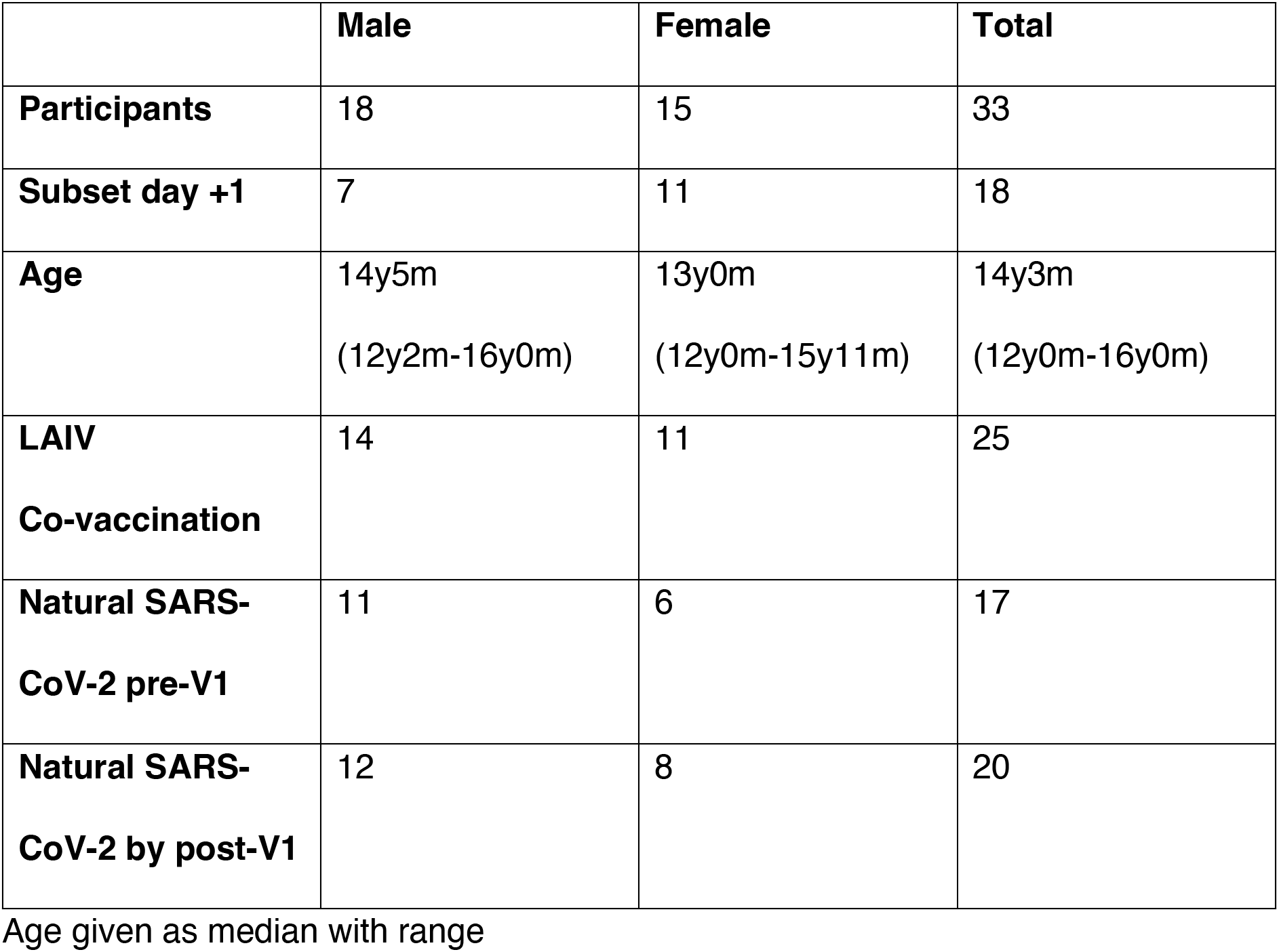
Cohort Demographics.

### High pDC activation and cytokine secretion following co-/vaccination

Since IFN-I plays a central role in the initiation of both innate and adaptive immune responses we first studied the behaviour of pDCs, as the main IFN-I producing cell type, to TLR7 in the 18 participants additionally sampled the day following co-/vaccination (Figure 1; see pDC Gating Strategy in Supplementary Figure S1).

**Figure 1–.**
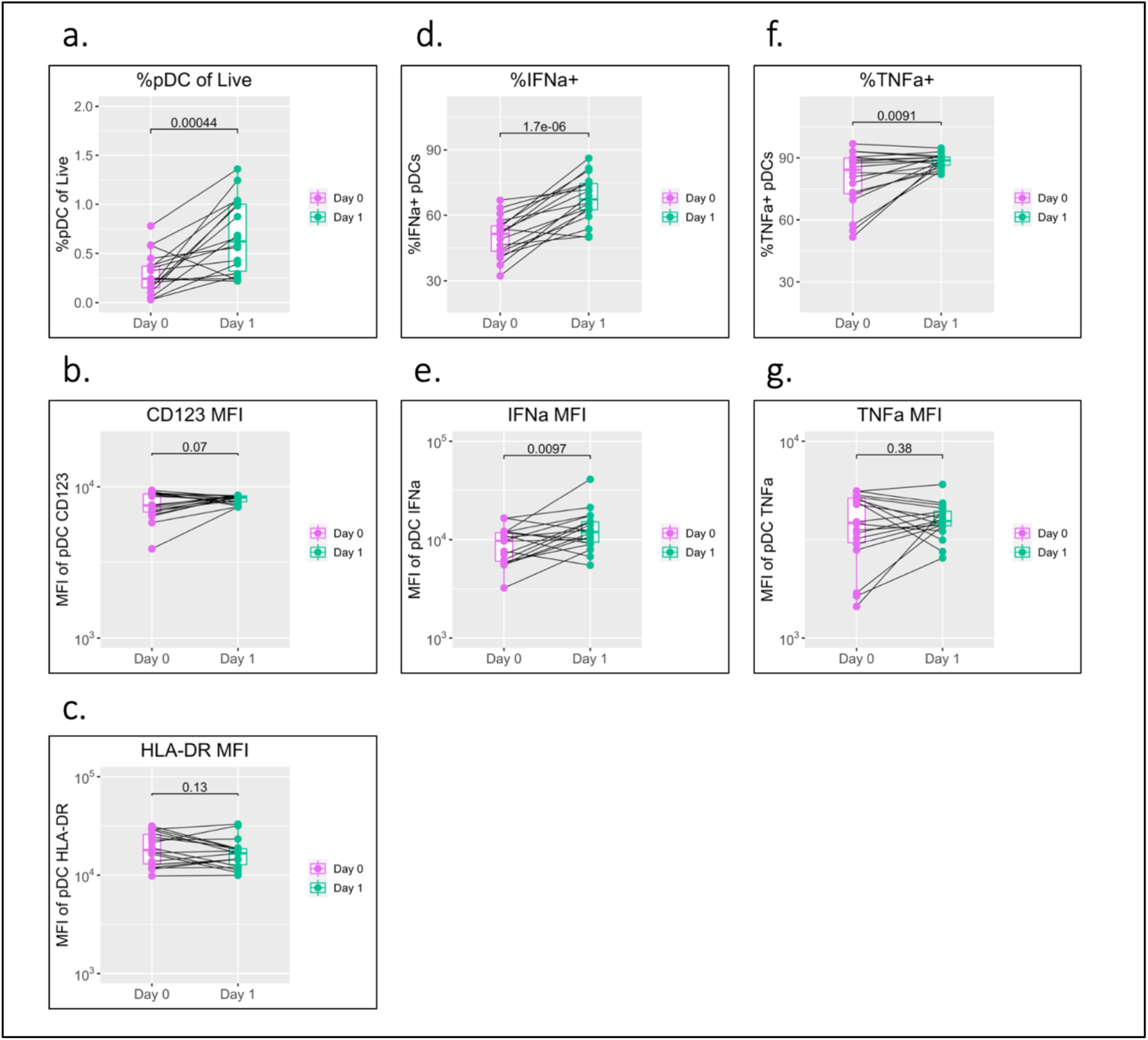
pDC activation following BNT162b2/LAIV co-/vaccination. pDC flow cytometry data following whole blood TLR7 stimulation for the day of vaccination (D0) and the following day (D1) (n=18; male=7, female=11. a) percentage of pDCs of live PBMC, b) MFI of pDC CD123, c) MFI of pDC HLA-DR, d) percentage of IFNa+ pDCs, e) MFI of pDC IFNa, f) percentage of pDC TNFa, g) MFI of pDC TNFa. Pairwise comparisons via paired Student’s T-test.

A striking increase in pDC abundance was observed between baseline (D0) and the following day (D1) with the pDC population as a percentage of total PBMC increasing from a median of 0.24% to 0.62% (p=0.00044) (Figure 1a.). Increased pDC activation between D0 and D1 was also seen in a modest increase in CD123 (p=0.07) but was not reflected in HLA-DR (Figure 1b. & c.).

pDC IFN-α production increased significantly between D0 and D1 with a significant increase in the percentage of IFN-α+ pDCs (from a median 51.5% to 67.3%; p<0.0001) and individual cell expression measured via MFI (median of 9694 to 11937; p=0.0097) (Figure 1d. & e.).

Production of TNF-α showed a mixed pattern in terms of the percentage of TNF-α+ pDCs and TNF-α MFI; levels tended to increase between D0 and D1 for individuals with low D0 expression and decrease for individuals with high D0 expression (Figure 1f. & g.).

pDC flow cytometry data collected for two unvaccinated control individuals showed consistency between D0 and D1 with unaltered MFI for CD123, HLA-DR and unaltered production of IFN-α and TNF-α (Figure S2).

### Delayed activation kinetics and IFN-I production in male pDCs

As described above, female pDCs produce greater quantities of IFN-I in response to TLR7 stimulation than male pDCs and we have recently reported that this may be underpinned by differential activation kinetics [22]. We therefore next stratified the data by sex to investigate whether differential activation kinetics were apparent between male and female pDCs following co-/vaccination (Figure 2).

**Figure 2–.**
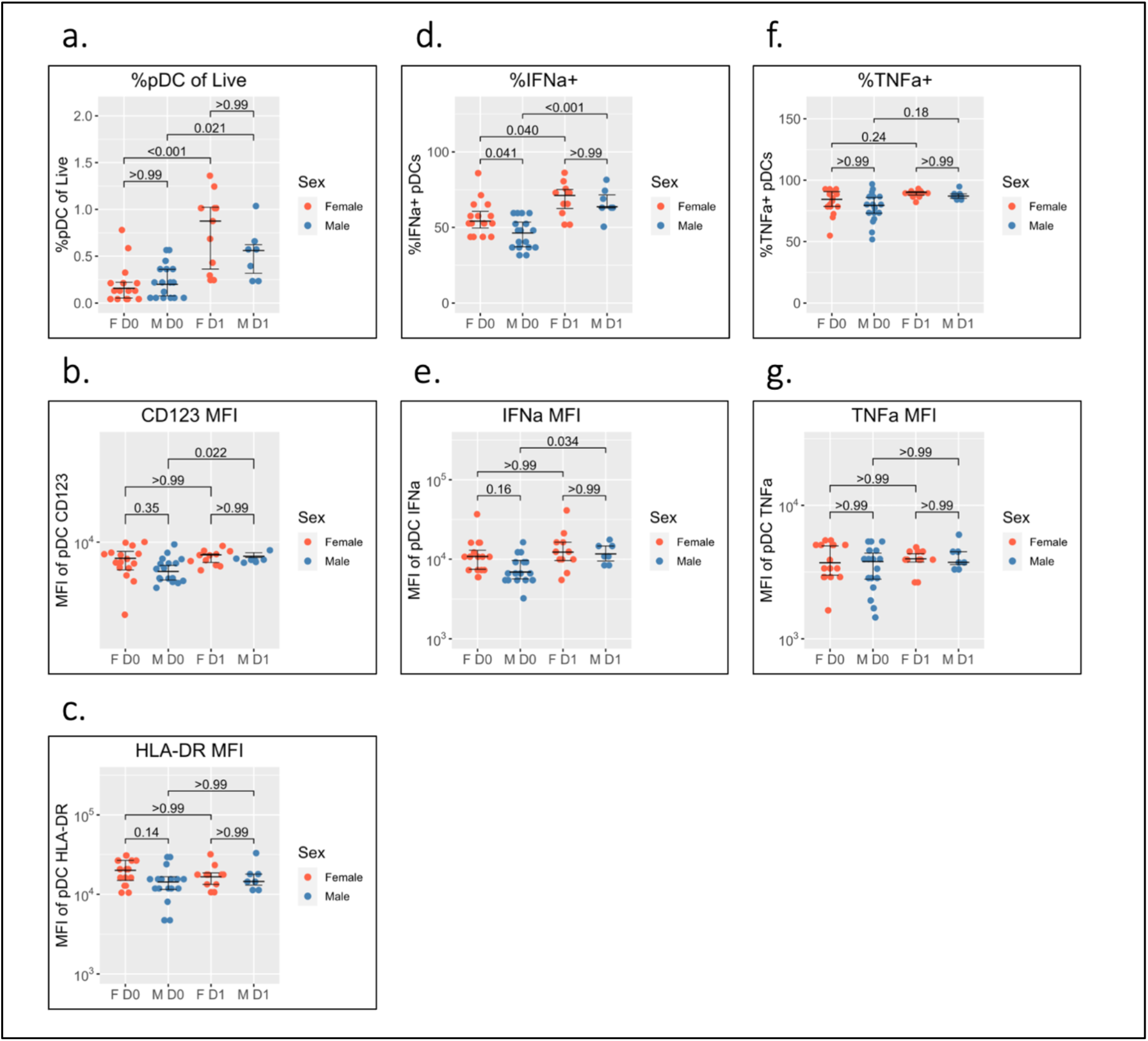
Sex differential pDC activation following BNT162b2/LAIV co-vaccination. Sex-stratified pDC flow cytometry data following whole blood TLR7 stimulation for the day of vaccination (D0) (n=33; male=18, female=15) and the following day (D1) (n=18; male=7, female=11). a) percentage of pDCs of live PBMC, b) MFI of pDC CD123, c) MFI of pDC HLA-DR, d) percentage of IFNa+ pDCs, e) MFI of pDC IFNa, f) percentage of pDC TNFa, g) MFI of pDC TNFa. Pairwise comparisons via unpaired Student’s T-test with Bonferroni correction for multiple comparisons.

pDC abundance is comparable between males and females on D0 and increases in both sexes to D1, but to a greater extent in females (median 0.16% to 0.88%; p<0.001) than males (median of 0.20% to 0.56%; p=0.021) (Figure 2a.).

pDC activation via CD123 appears lower on D0 for males than females but does not reach statistical significance (p=0.35). There is a significant increase in CD123 expression between D0 and D1 for male pDCs (5204 to 6469; p=0.022) whereas female CD123 remains consistent such that no difference is apparent between the sexes on D1 (Figure 2b.). A similar pattern was also observed for HLA-DR where expression appeared higher for female pDCs on D0 but was comparable to male pDCs on D1 (Figure 2c.).

The widely reported significantly increased IFN-α production by female pDCs is also seen here on D0 (median 54.1% compared to 46.3% for males; p=0.041) (Figure 1d.). The percentage of IFN-α+ pDCs increases for both sexes between D0 and D1 but more so for male pDCs (46.3% to 63.6%; p<0.001) than females (54.1% to 71.2%; p=0.040) such that there is no statistical sex difference on D1 (Figure 2d.). The same pattern is seen for IFN-α MFI (Figure 2e.).

Conversely, no sex difference is seen for the percentage of TNF-α+ pDCs nor TNF-α MFI either before or after co-/vaccination (Figure 2f. & g.).

### Gonadal androgens but not adrenal androgens differ between males and females and correlate with age during adolescence

The pubertal age range of the adolescents studied offered a unique opportunity to investigate further the relationship between androgens and pDC IFN-I production. In addition to canonical gonadal androgen synthesis – which produces testosterone and dihydrotestosterone (DHT) via the precursor androstenedione – adrenal synthesis of 11-oxygenated androgens is reported to contribute significantly to the circulating androgen pool. Serum levels of both gonadal and adrenal androgens were therefore determined for each participant via liquid chromatography tandem mass spectrometry (Figure 3). Since nearly all circulating testosterone is bound by either sex hormone binding globulin (SHBG) or albumin and is biologically unavailable to cells in the blood [33], the level of unbound, or ‘free’, testosterone was also calculated using measurements of SHBG and albumin [34].

**Figure 3–.**
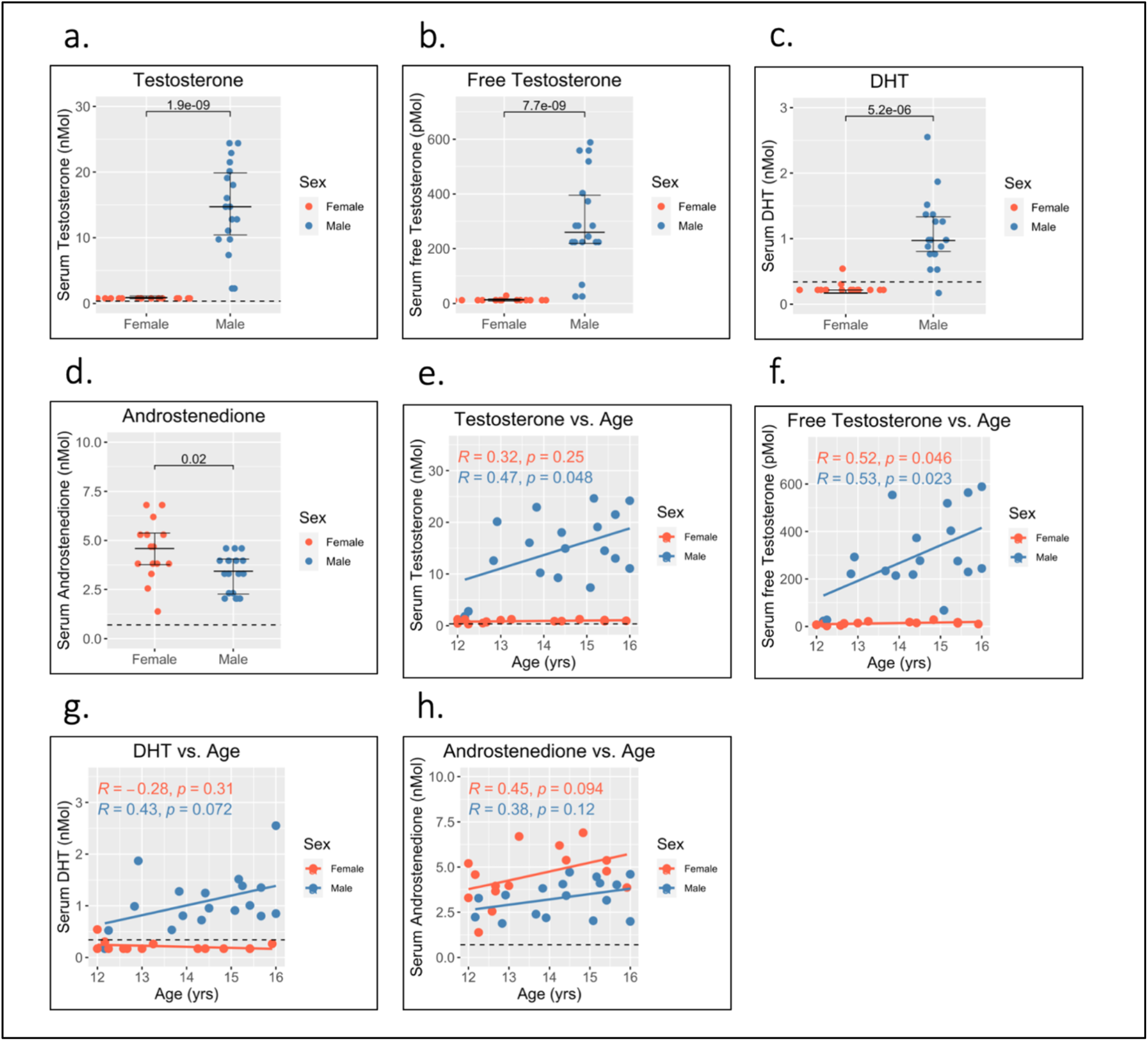
Serum gonadal androgen measurements in adolescent males and females. Serum androgen concentrations determined via tandem mass spectrometry for adolescents (n=33; male=18, female=15). a) to d) comparison of serum concentrations in males and females for: testosterone, free testosterone, dihydrotestosterone (DHT), and androstenedione, respectively. e) to h) correlation of androgen concentration with age for males and females for: testosterone, free testosterone, dihydrotestosterone (DHT), and androstenedione, respectively. Pairwise comparisons via Student’s unpaired t-test. Correlations via Pearson’s method.

As expected, levels of testosterone, free testosterone, and DHT were all significantly higher for male adolescents than female adolescents (Figure 3a. to c.) and all increased with age in males (Figure 3e. to g.). Conversely, levels of androstenedione were significantly higher for females (Figure 3d.), and weakly correlated with age in females but not males (Figure 3h.).

Levels of 11-oxygenated androgens and DHEA were not significantly different between males and females (Figure 4a. to e.) which suggests that adrenal androgens do not contribute significantly to sex differences in androgen levels. Further 11-OH androstenedione (11OHA4) was the only adrenal androgen to correlate with age (Figure 4f. to j.) and increased with age in females (Figure 4g.).

**Figure 4–.**
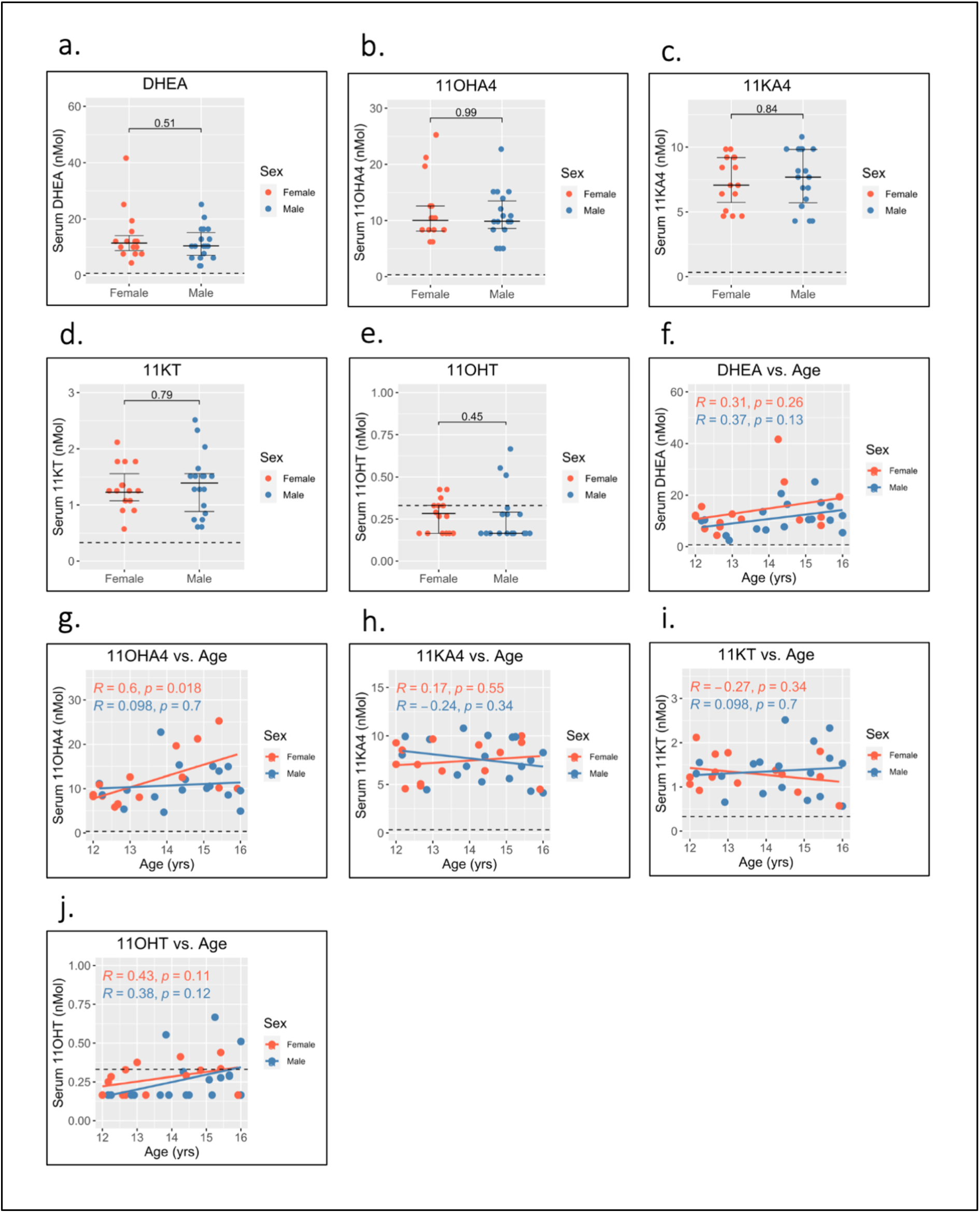
Serum adrenal androgen measurements in adolescent males and females. Serum androgen concentrations determined via tandem mass spectrometry for adolescents (n=33; male=18, female=15). a) to e) comparison of serum concentrations in males and females for: dehydroepiandrosterone (DHEA), 11-hydroxyandrostenedione (11OHA4), 11-ketoandrostenedione (11KA4), 11-ketotestosterone (11KT), 11-hydroxytestosterone (11OHT), respectively. f) to j) correlation of androgen concentration with age for males and females for: dehydroepiandrosterone (DHEA), 11-hydroxyandrostenedione (11OHA4), 11-ketoandrostenedione (11KA4), 11-ketotestosterone (11KT), 11-hydroxytestosterone (11OHT), respectively. Pairwise comparisons via Student’s unpaired t-test. Correlations via Pearson’s method.

### pDC IFN-I production is negatively associated with free testosterone

Since androgens have been implicated in a dampened vaccine response [10] and in reduced IFN-I production in male infants [25] we next sought to investigate whether the androgen levels measured here related to pDC IFN-I production. Using D0 flow cytometry data to control for any effect of co-/vaccination, a Least Absolute Shrinkage and Selection Operator (LASSO) regression model [35,36] was constructed using androgen concentrations as input co-variates and the percentage of IFN-α+ pDCs as the dependent variable to determine if a relationship exists between androgens and pDC IFN-α+ (Table 2).

**Table 2–.**
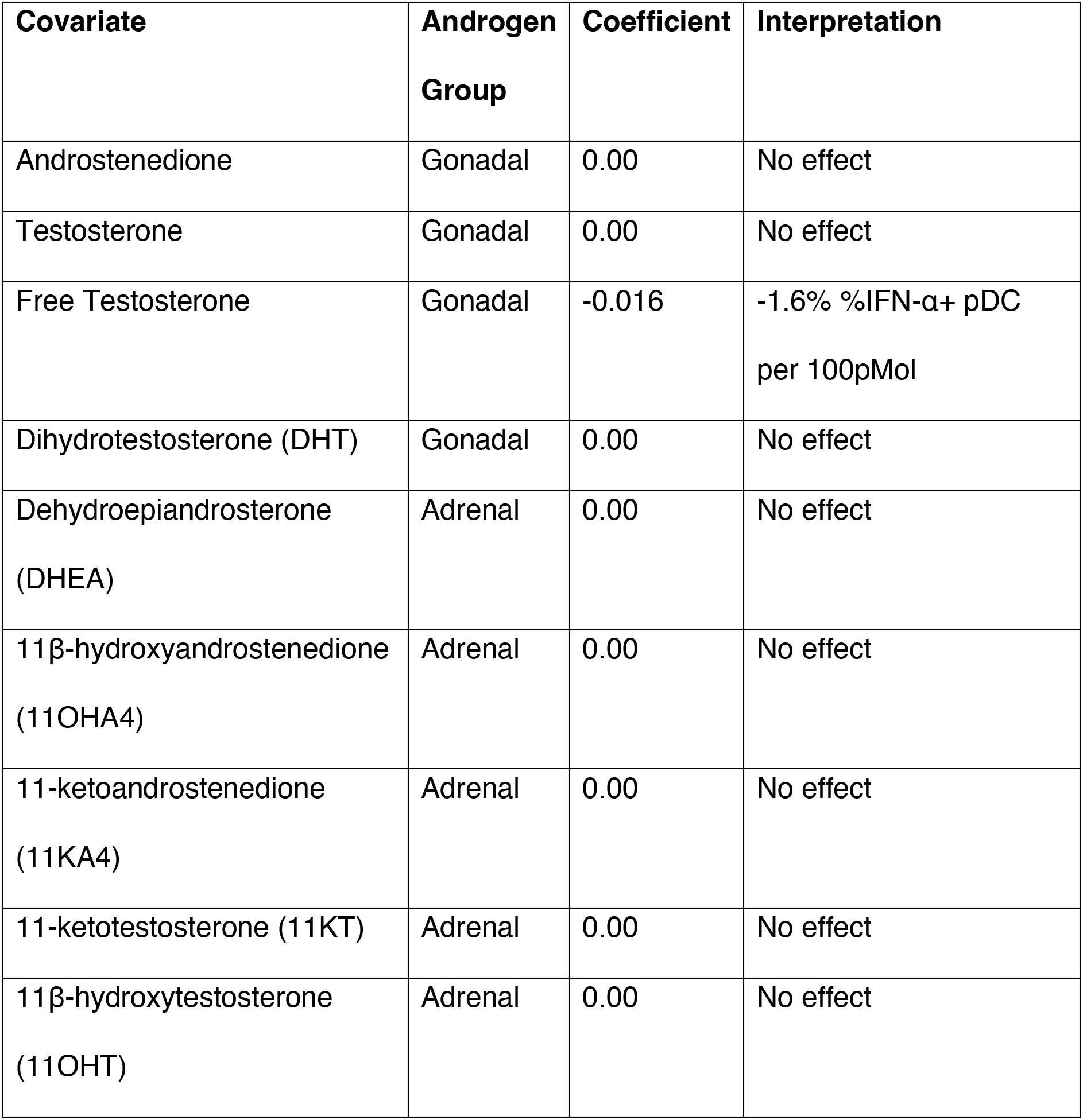
LASSO regression co-efficient estimates for the effect of androgens on pDC IFN-I.

Only free testosterone is identified by the model as influential of pDC IFN-α and causes a reduction of 1.6% IFN-α+ pDCs per 100pMol. Moreover, this effect overshadowed a negative influence by total testosterone which was only identified if free testosterone was excluded from the model (Table S1).

### BNT162b2 IgG response is increased by LAIV co-vaccination and DHT but decreased by pDC abundance

Given the potential of IFN-I to influence each stage of the immune response to vaccination we next turned to the relationship between the early pDC IFN-I response and the outcome of BNT162b2 vaccination. Having observed differential activation between male and female pDCs (Figure 2) and having identified a role for free testosterone in decreased pDC IFN-I (Table 2) we analysed the D0 flow cytometry data together with gonadal androgen measurements in LASSO models for post-V1 anti-Spike-IgG and anti-RBD-IgG responses (Table 3). After controlling for natural SARS-CoV-2 infection, LASSO modelling was only possible for infected adolescents (n=20) due to the small number of uninfected adolescents (n=13).

**Table 3–.** LASSO regression co-efficient estimates for post-V1 anti-Spike and anti-RBD IgG.

| Covariate | Spike<br>Coefficient | Interpretation | RBD<br>Coefficient | Interpretation |
| --- | --- | --- | --- | --- |
| Female Sex | 0.00 | No effect | 0.00 | No effect |
| Age | 0.00 | No effect | 0.00 | No effect |
| LAIV Co-vaccination | 221,683 | +2.2x10<sup>5</sup> AU<br>if co-vaccinated | 91,350 | +9.1x10<sup>4</sup> AU<br>if co-vaccinated |
| %pDC/Live | -83,379 | -8.3x10<sup>3</sup> AU<br>per extra<br>0.1% pDC | -26,624 | -2.6x10<sup>3</sup> AU<br>per extra<br>0.1% pDC |
| CD123 MFI | 0.00 | No effect | 0.00 | No effect |
| HLA-DR MFI | 0.00 | No effect | 0.00 | No effect |
| %TNF-α+ | 0.00 | No effect | 0.00 | No effect |
| %IFN-α+ | 0.00 | No effect | 0.00 | No effect |
| Androstenedione | 0.00 | No effect | 0.00 | No effect |
| Testosterone | 0.00 | No effect | 0.00 | No effect |
| Free<br>Testosterone | 0.00 | No effect | 0.00 | No effect |
| DHT | 65,047 | +6.5x10<sup>4</sup> AU<br>per nMol DHT | 90,845 | +9.1x10<sup>4</sup> AU<br>per nMol DHT |

The IgG titres to BNT162b2 are reported elsewhere [32] and were significantly higher in males than females for SARS-CoV-2 naïve adolescents although they did not differ between the sexes for infected adolescents. Likewise, there was no sex difference in the IgG titres of the four LAIV haemagglutinin antigens [32].

Surprisingly, the largest effect is seen for LAIV co-vaccination which is associated with increased titres for both anti-spike-IgG and anti-RBD-IgG. In contrast to reports of androgens suppressing vaccine responses, DHT is also identified as positively influencing IgG titres for both antigens. The only other association identified by the model is a negative relationship between pDC abundance and the IgG titres of both antigens.

The size of the model (n=20) precluded the inclusion of adrenal androgens but these were found not to influence IgG titres to either antigen in separate LASSO analyses containing only gonadal and adrenal androgens (Table S2).

## Discussion

This study sought to characterise the behaviour of pDCs from adolescents in response to BNT162b2/LAIV co-/vaccination and investigate whether a role can be attributed to differential pDC IFN-I in the widely reported sex differential outcome to vaccination, particularly to seasonal TIV [5,7,8,9,10,11,37].

After BNT162b2/LAIV co-/vaccination in adolescents a substantial increase in pDC abundance was observed by the following day, as was a modest increase in cell activation measured via CD123 expression. This accompanied a larger increase in cytokine production as both IFN-α and TNF-α were significantly increased the day following vaccination.

Importantly, differences were seen in the behaviour of male and female pDCs. On the day of vaccination male pDCs produced significantly less IFN-α than female pDCs in line with previous studies in children and adults [15,16,17,18,19,20,21,22]. Male pDC IFN-α increased to a greater extent than for female pDCs the day following vaccination, and was accompanied by an increase in cell activation by greater CD123 expression, whilst the abundance of pDCs increased more so for females than males. This aligns with our previous observations that biallelic TLR7 expression may allow female pDCs to more readily produce IFN-I whilst male pDCs require a greater level of activation in order to catch up [22]. Indeed, higher levels of the T-cell co-stimulatory molecule CD86 have been reported on female pDCs compared to male pDCs 20hrs following TLR7 stimulation [20]. Such need for increased activation by male cells may provide one explanation for the greatest risk group for adverse effects following SARS-CoV-2 mRNA vaccination being young males [38].

In addition, the pubertal age range of the cohort gave the unique opportunity to investigate whether the immunosuppressive properties of androgens extend to pDC IFN-I in adolescents. Males had significantly higher levels of the classic gonadal androgens but despite reports of their contribution to the circulating androgen pool 11-oxygenated adrenal androgens were found not to differ significantly between males and females. Indeed LASSO modelling using the hormone data identified only free testosterone as negatively influential of pDC IFN-α. That this effect overshadowed the effect of total testosterone implies a direct antagonistic relationship between testosterone and pDC IFN-α as nearly all circulating testosterone is protein-bound [33] and therefore unavailable to act on cells within the blood.

IFN-I is important for priming an adaptive immune response by activating CD4+ T-helper cells and B-cells [3], and the female adaptive immune system is predominantly T_H_2-biased [11]. It might therefore be anticipated from our data showing increased female pDC IFN-I following BNT162b2/LAIV co-/vaccination that anti-spike-IgG and anti-RBD-IgG antibody titres are also increased in females. Conversely, titres for both are higher in males for SARS-CoV-2 naïve adolescents, and equal for males and females after SARS-CoV-2 infection [32]. Unsurprisingly then we did not identify sex or any measure of pDC activation and cytokine production, including free testosterone, in LASSO models for anti-Spike-IgG and anti-RBD-IgG using data from SARS-COV-2 infected adolescents. Instead, LAIV co-vaccination was identified as the strongest positive influence of antibody titres. Somewhat more surprisingly DHT was also identified as positively influencing antibody titres to both antigens.

A similar difference was also seen by Sparks *et al*. [39] who reported increased antibody titres to the quadrivalent seasonal influenza vaccine in males compared to females following natural SARS-CoV-2 infection and link this to an increase in “virtual memory” T-cells in males. Such non-specific activation of T-cells unrelated to the present vaccine immune response, so-called “bystander activation”, has also been reported for Modified Vaccinia Ankara (MVA) [40] and Bacillus Calmette-Guerin (BCG) [41], and has been attributed to IFN-I induced by vaccines [42]. LAIV is a potent inducer of IFN-I [43] and IFN-I signatures 7 days post-vaccination have previously been shown to be predictive of neutralising antibody titres to LAIV [6]. Importantly this was attributable in children to increased numbers of total naïve, memory, and transitional B-cells following LAIV vaccination when compared to seasonal TIV [6], which is a weaker inducer of IFN-I relative to LAIV [43].

It follows that the boosting effect of LAIV seen here may be attributable to increased levels of systemic IFN-I leading to non-specific bystander activation of SARS-CoV-2 memory cells. That IFN-I is not directly identified in the LASSO models for anti-spike-IgG or anti-RBD-IgG may be explained by the fact that LAIV IFN-I production does not peak until 7 days post-vaccination whereas the pDC data used in the LASSO models was from the day of vaccination.

The main limitation of our study is that number of adolescents studied was lower than planned due to reduced recruitment resulting from the SARS-CoV-2 Delta wave in autumn 2021 [44]. This limited the LASSO analyses that could be performed and reduced the power of the study to identify effects that were statistically significant.

The IFN-I elicited by LAIV can be attributed to TLR7 signalling [14,43] and whilst IFN-I is a powerful vaccine adjuvant, excessive induction leads to hyperinflammation and autoimmunity [45]. Accordingly, mRNA vaccines like BNT162b2 are formulated to minimise TLR7 signalling via highly efficient purification through high-pressure liquid chromatography and substitution for the hypo-agonistic synthetic nucleotide 1-methylpseudouridine [46,47,48]. mRNA preparations with such modifications display an extended cytoplasmic half-life and translational potential *in-vitro* [49,50,51]. It follows that TLR7 activation may therefore increase the rate of clearance of mRNA within cells and decrease the cytoplasmic half-life of mRNA vaccines *in-vivo*. Although present in circulation, DHT is mainly synthesised locally by specialised target tissues including the skin [52] and its identification as positively influential of BNT162b2 antibody titres suggests that DHT could act locally to reduce TLR7 signalling within vaccination sites to increase the duration of antigen availability for the humoral immune response.

This deviates from the current understanding of the immune-suppressive role of androgens in reducing vaccine responses in males compared to females [10,12,23]. However, BNT162b2 antibody titres were higher in males than females for both antigens in SARS-CoV-2 naïve adolescents and were comparable for males and females after natural SARS-CoV-2 infection in this cohort [32]. Likewise, no sex difference was seen for LAIV [32], and has not been reported to-date in other studies of LAIV. LAIV replicates within cells of the upper airway mucosa [53] and BNT162b2 is translated within muscle cells [48]. Together these data suggest that the influence of androgens on vaccines with intracellular stages is different from the immunosuppressive role reported for seasonal protein-subunit influenza vaccines. Local modulation of the IFN-I response by androgens may therefore be beneficial for the outcome of such vaccines.

This study highlights the role played by IFN-I in the outcome to BNT162b2 + LAIV co-/vaccination and identified sex differences in the primary IFN-I producing cell type, pDCs. Free testosterone was seen to reduce pDC IFN-I but DHT was associated with increased BNT162b2 antibody titres. As understanding of the molecular triggering events surrounding empirically developed vaccine adjuvant strategies increases, so will the need to greater understand the complex role played by differential early IFN-I responses in determining the outcome of new vaccines.

## Materials and Methods

### Ethics Statement

Blood was sampled from adolescent participants in accordance with approval from the University of Oxford Medical Sciences Interdivisional Research Ethics Committee (reference R71346/RE001). Written, informed parental consent with individual assent was obtained for each participant.

### Whole blood measurement of pDC IFN-I

pDC IFN-I was measured via whole blood stimulation as previously published [22]. Briefly, blood was collected immediately after vaccination directly into heparin vacutainers (BD) containing R10 + Brefeldin A + CL097 and maintained at 37C in a dry block-heater during laboratory transport before further incubation up to 6hrs, RBC lysis, and staining for flow cytometry via LSR II.

### Determination of serum androgen concentrations

Blood was collected concurrently at the time of vaccination and at a median of 37 days post-vaccination (“post-V1”) via SST II Advance tubes (BD). Samples were Serum was isolated via centrifugation 2000g 7mins and stored at −80C. Steroids were extracted from 200μl of serum and quantified using liquid chromatography tandem mass spectrometry, as described previously [54,55]. Values below the lower limit of qualification (LLOQ) were set to half the LLOQ. The average of measurements for the day of vaccination and post-V1 was used for analysis.

The concentration of serum free testosterone was calculated via the Vermeulen equation [34] using the concentration of serum sex hormone binding globulin and albumin determined using a Cobas C system (Roche) via electrochemiluminescence (Elecsys SHBG) and bromocresol purple assay (ALBP), respectively. The average of measurements for the day of vaccination and post-V1 was used for analysis.

### Determination of anti-SARS-CoV-2 IgG responses and SARS-CoV-2 infection

Blood was collected concurrently on the day of vaccination and post-V1 via EDTA vacutainers (BD) and plasma isolated by Ficoll (Lymphoprep) separation (1000g; 15mins; low deceleration) and stored at −80C. IgG responses to SARS-CoV-2 Spike, receptor-binding domain (RBD), and nucleocapsid (N) were determined via Meso Scale Discovery (MSD) V-plex immunoassay ‘Coronavirus Panel 3’, as described previously [56].

SARS-CoV-2 infection was defined using the MSD cut-offs determined previously [56]. Prior infection was defined as either anti-Spike-IgG or anti-N-IgG above the cutoff and infection post-V1 defined as anti-N-IgG above the cutoff due to the inclusion of Spike in BNT162b2.

### Data Analysis and LASSO modelling

Flow cytometry data was analysed with FlowJo v10.7.1 (BD) and pDCs defined as: PBMC/Singlet/Live/CD56-/CD19-/CD3-/CD14-/CD11c-/CD123+/HLA-DR+ [22].

Graphs were plotted and analysed via R Studio running R v4.3.0 via base R [57] and package *ggpubr*. Two-group continuous variables were analysed via Student’s t-test (parametric data) and Wilcoxon test (non-parametric data). Were appropriate, Welch’s correction was applied for unequal variances and Bonferoni correction used to correct multiple comparisons.

The Least Absolute Shrinkage and Selection Operator (LASSO) regression model assigns a penalty on the absolute value of each covariate’s regression coefficient and, hence, shrinks them towards zero, setting irrelevant variables to exactly zero. This way, variable selection is achieved and any coefficients not regressed to zero therefore have an effect on the dependent variable in the model [35]. The R package *glmnet* [36] was used to create relaxed-fit LASSO models using *lambda.min* from model cross-validation.

### Data Availability Statement

Analyses in this manuscript combine only raw data presented in this manuscript with raw data published elsewhere (Jay et al. (2023) medRxiv).

## Supporting Information

**Figure S1–.**
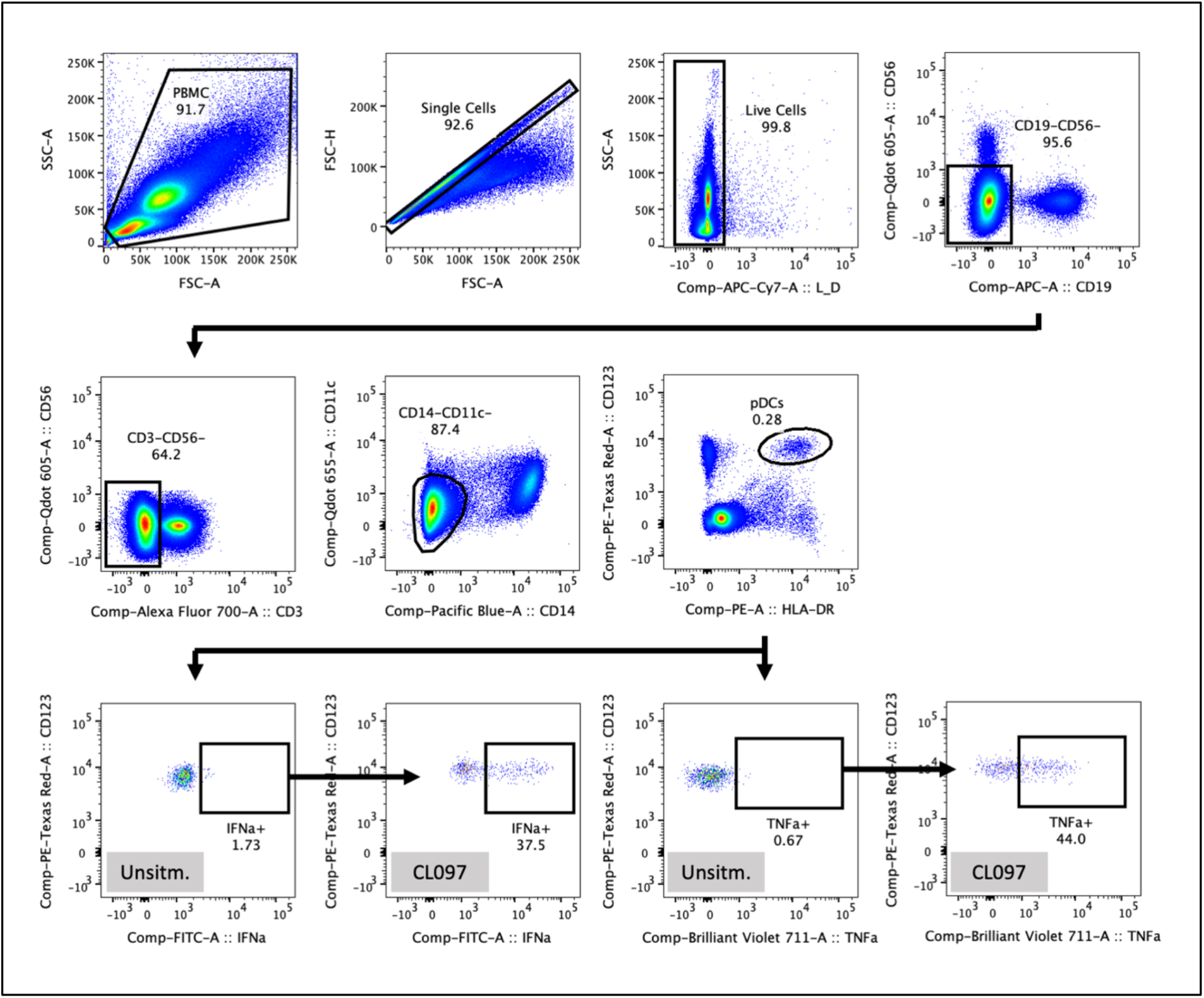
pDC Gating Strategy. Representative gating strategy for isolating pDCs from lysed whole blood. After PBMC, Singlet, and Live Cell gating, pDCs were defined as: CD19-/CD56-/CD3-/CD14-/CD11c-/HLA-DR+/CD123+. Cytokine gates for IFNa and TNFa were set using unstimulated controls and applied to CL097-stimulated samples, as shown.

**Figure S2–.**
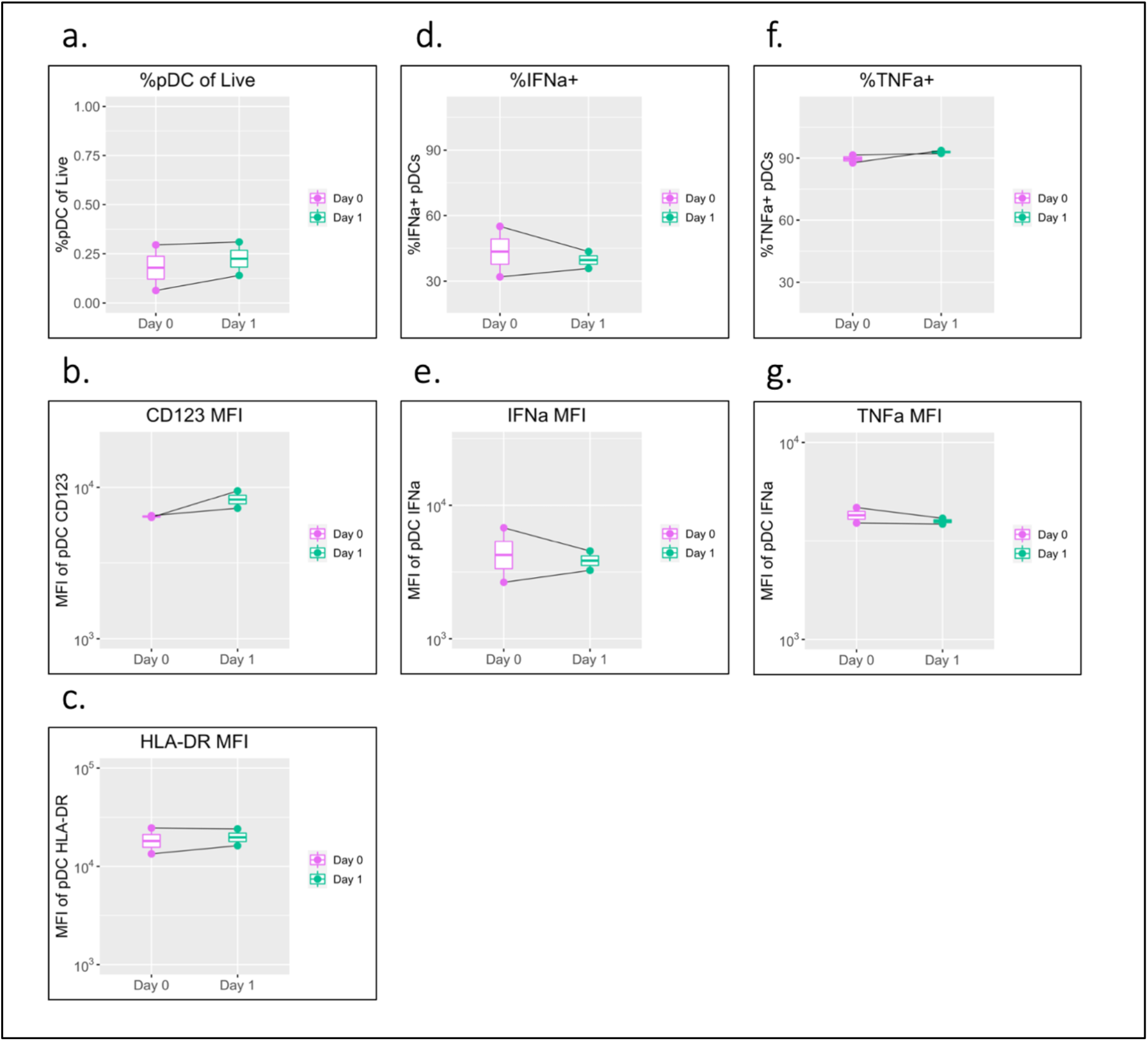
pDC phenotype of unvaccinated control individuals. pDC flow cytometry data following whole blood TLR7 stimulation concurrently collected from two unvaccinated control individuals on the day of vaccination (D0) and the following day (D1). a) percentage of pDCs of live PBMC, b) MFI of pDC CD123, c) MFI of pDC HLA-DR, d) percentage of IFNa+ pDCs, e) MFI of pDC IFNa, f) percentage of pDC TNFa, g) MFI of pDC TNFa. Pairwise comparisons via paired Student’s T-test.

**Table S1–.** LASSO regression co-efficient estimates for the effect of androgens on pDC IFN-I excluding free testosterone.

| Covariate | Androgen Group | Coefficient | Interpretation |
| --- | --- | --- | --- |
| Androstenedione | Gonadal | 0.00 | No effect |
| Testosterone | Gonadal | -0.32 | -0.32% IFN-α+ per nMol |
| Dihydrotestosterone (DHT) | Gonadal | 0.00 | No effect |
| Dehydroepiandrosterone (DHEA) | Adrenal | 0.00 | No effect |
| 11β-hydroxyandrostenedione (11OHA4) | Adrenal | 0.00 | No effect |
| 11-ketoandrostenedione (11KA4) | Adrenal | 0.00 | No effect |
| 11-ketotestosterone (11KT) | Adrenal | 0.00 | No effect |
| 11β-hydroxytestosterone (11OHT) | Adrenal | 0.00 | No effect |

**Table S2–.**
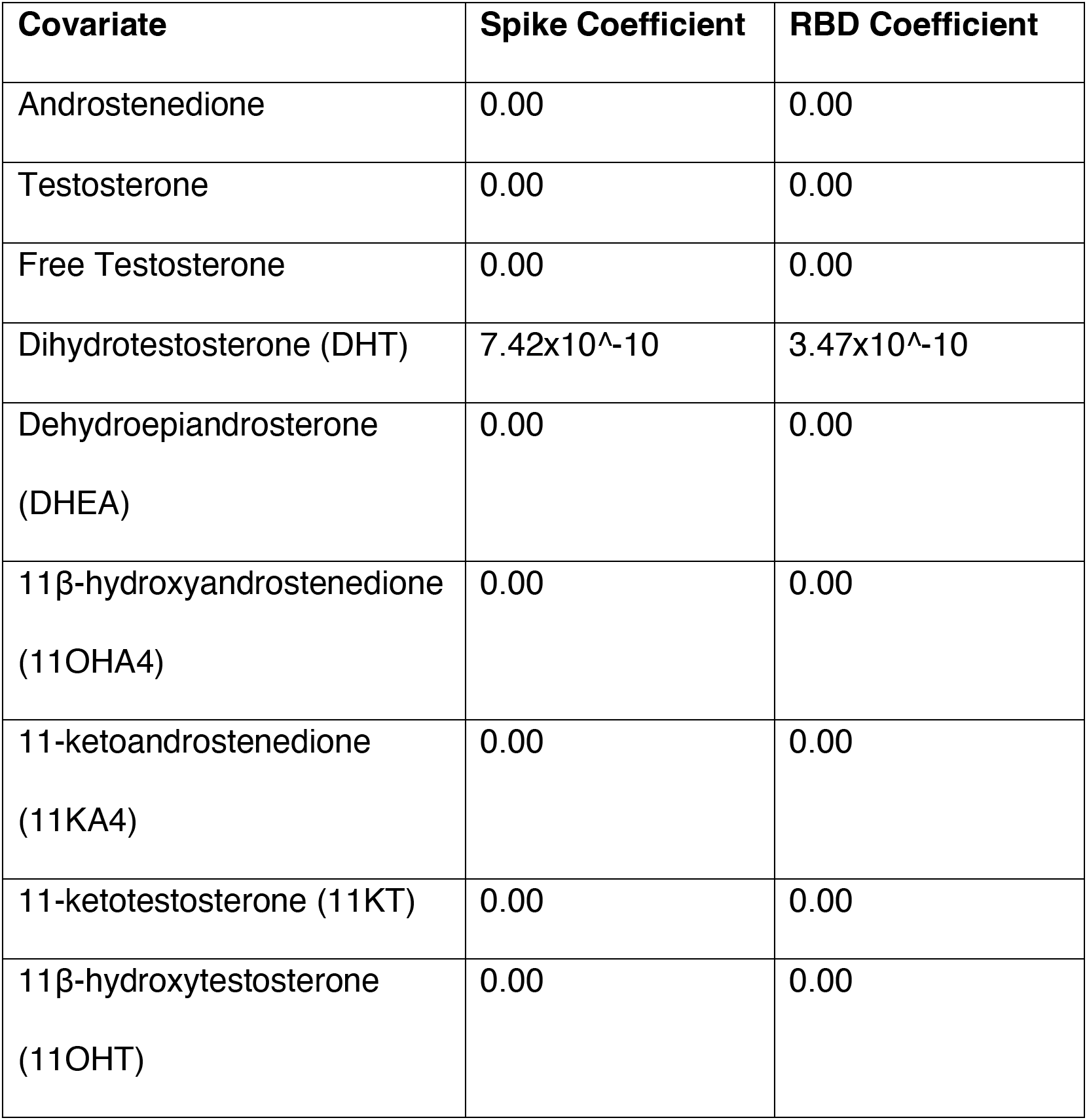
LASSO regression co-efficient estimates for the effect of androgens on post-V1 anti-Spike and anti-RBD IgG.

| Covariate | Spike Coefficient | RBD Coefficient |
| --- | --- | --- |
| Androstenedione | 0.00 | 0.00 |
| Testosterone | 0.00 | 0.00 |
| Free Testosterone | 0.00 | 0.00 |
| Dihydrotestosterone (DHT) | $7.42 \times 10^{-10}$ | $3.47 \times 10^{-10}$ |
| Dehydroepiandrosterone<br>(DHEA) | 0.00 | 0.00 |
| 11β-hydroxyandrostenedione<br>(11OHA4) | 0.00 | 0.00 |
| 11-ketoandrostenedione<br>(11KA4) | 0.00 | 0.00 |
| 11-ketotestosterone (11KT) | 0.00 | 0.00 |
| 11β-hydroxytestosterone<br>(11OHT) | 0.00 | 0.00 |

